# Medial prefrontal cortex suppresses reward-seeking behavior with risk of positive punishment by reducing sensitivity to reward

**DOI:** 10.1101/2024.01.03.573768

**Authors:** Monami Nishio, Masashi Kondo, Eriko Yoshida, Masanori Matsuzaki

**Author notes:** Correspondence (M.M.) Address: Department of Physiology, Graduate School of Medicine, The University of Tokyo, 7-3-1, Hongo Bunkyo-ku, Tokyo 113-0033, Japan. These authors contributed equally (Monami Nishio, Masashi Kondo).

## Abstract

Reward-seeking behavior is frequently associated with risk of punishment. There are two types of punishment: positive, resulting in an unpleasant outcome, and negative, resulting in omission of a reinforcing outcome. Although the medial prefrontal cortex (mPFC) is important in avoiding punishment, whether it is important for avoiding both positive and negative punishment and how it contributes to such avoidance are not clear. In this study, we trained male mice to perform decision-making tasks under the risks of positive (air-puff stimulus) and negative (reward omission) punishment. We found that pharmacological inactivation of mPFC enhanced the reward-seeking choice under the risk of positive, but not negative, punishment. In reinforcement learning models, this behavioral change was well-explained by hypersensitivity to the reward, rather than a decrease in the strength of aversion to punishment. Our results suggest that mPFC suppresses reward-seeking behavior by reducing sensitivity to reward under the risk of positive punishment.

## Introduction

You know sweets are probably there. If you find them and eat them, your mother might get mad at you. Will you go there? Actions motivated by rewards are frequently associated with the risk of punishment^1^. The risk of an adverse consequence resulting from a reward-seeking behavior can profoundly affect subsequent behavior. Such adverse events can be described as punishment because they reduce the probability that the same behavior will be produced again and increase the exploration of less risky alternatives. Punishment can be thought of as being either positive or negative, with positive punishment resulting in directly experiencing an unpleasant outcome (e.g., a stimulus that causes discomfort or pain) and negative punishment omitting a reinforcing consequence (e.g., risk of loss or reinforcer uncertainty)^2–4^. Avoiding the negative punishment is synonymous with obtaining a covert reward (which could serve as a reinforcer of “doing nothing” in the choice of to perform an action or not)^5^. The brain must compute the threat of punishment for the optimal selection and execution of actions. This ability is known to be impaired in a variety of neuropsychiatric disorders, such as addiction, which is characterized by the persistence of behaviors with the risk of short- and long-term adverse consequences^4,6,7,8^.

It is well known that the medial prefrontal cortex (mPFC) is involved in decision-making during approach-avoidance conflicts^9–18^. A strong response of the dorsomedial prefrontal cortex (dmPFC) to risk is associated with less risky choice by humans^9^, and patients with a ventromedial prefrontal cortex (vmPFC) lesion can show hypersensitivity to reward^10^. Many neurons in the macaque anterior cingulate cortex (ACC) represent the value and uncertainty of rewards and punishments^11^. In rats, the silencing of mPFC neurons promotes cocaine-seeking behavior under the risk of foot shock^12^, and binge drinking of alcohol under the risk of a bitter tastant (quinine)^13^, whereas activating the mPFC can attenuate these behaviors. mPFC neurons that project to the nucleus accumbens are suppressed prior to reward-seeking behavior with foot-shock punishment, while their activation reduces reward seeking^14^. Thus, mPFC is obviously involved in decision-making under a risk of positive punishment.

There are also a few studies that have examined the functions of mPFC under risk of negative punishment. For example, mPFC is known to be required for updating the choice bias in a choice task between a small reward with certainty and a large reward with high omission probability^4,19^. However, it has not been examined whether mPFC is equally involved in decision-making with positive and negative punishments in very similar tasks. In addition, it remains unclear which intrinsic variables mPFC regulates to avoid positive or negative punishment.

To address these issues, we examined the effect of pharmacological inactivation of mPFC on the probability of performing a lever pull that yields a reward under the risk of positive (air-puff stimulus) and negative (reward omission) punishments in head-fixed male mice. Although the head-fixed condition was more stressful for the mice than a free-moving condition, we head-fixed them using a method developed in a previous study^5^ so that one-photon and two-photon calcium imaging could be used to detect the relevant cortical activity in future experiments^20-22^. We found that mPFC is necessary for behavioral inhibition under the risk of positive punishment, but not negative punishment. Furthermore, we introduced an analytical method to explain the behavioral effect of mPFC inactivation in the form of a change of parameters in the context of reinforcement learning. Our analyses suggest that the effect of mPFC inactivation was best explained as an increase in the sensitivity to reward (that is, hypersensitivity to reward), rather than as a decrease in the strength of the aversion to positive punishment or a decrease in the covert reward to non-action.

## Results

### Both positive and negative punishments suppress the reward-seeking behavior of mice

We modified a two-tone lever-pull task for head-fixed mice that we previously reported^5^ into a two-tone lever-pull task with the risk of two types of punishment: positive punishment (i.e., exposure to an air-puff) and negative punishment (i.e., omission of a reward) (Figures 1A and 1B). Two pure tones (6 and 10 kHz pure tone for 0.8–1.2 s), the reward-seeking action (pulling the lever after the go sound cue [pink noise] that followed the tone presentation), and the reward (a water drop delivered from a spout near the mouth) were common to both tasks. In the two-tone lever-pull task with the risk of an air-puff (“air-puff task”), if the head-fixed mouse pulled the lever after the go cue presentation, a water reward was delivered at a probability of 100% regardless of the tone type. In addition, an air-puff (0.3–0.4 Mpa) was delivered at a probability of 90% after the lever was pulled in each trial with a tone A presentation (tone A trial) and a probability of 10% after the lever was pulled in each trial with a tone B presentation (tone B trial). Tones A and B were randomly presented at probabilities of 20% and 80%, respectively. If the animal did not pull the lever, they did not receive the air-puff or the water reward (Figure 1C). In the two-tone lever-pull task with a risk of reward omission (“omission task”), the water reward was delivered at a probability of 10% after a lever-pull in each tone A trial and at a probability of 90% after a lever-pull in each tone B trial. This corresponded to reward omission probabilities of 90% and 10% for tone A and B trials, respectively. Tones A and B were randomly represented at probabilities of 40% and 60%, respectively. No air-puff was delivered in the omission task, and if the mice did not pull the lever, they did not receive the water reward (Figure 1D).

**Figure 1.**
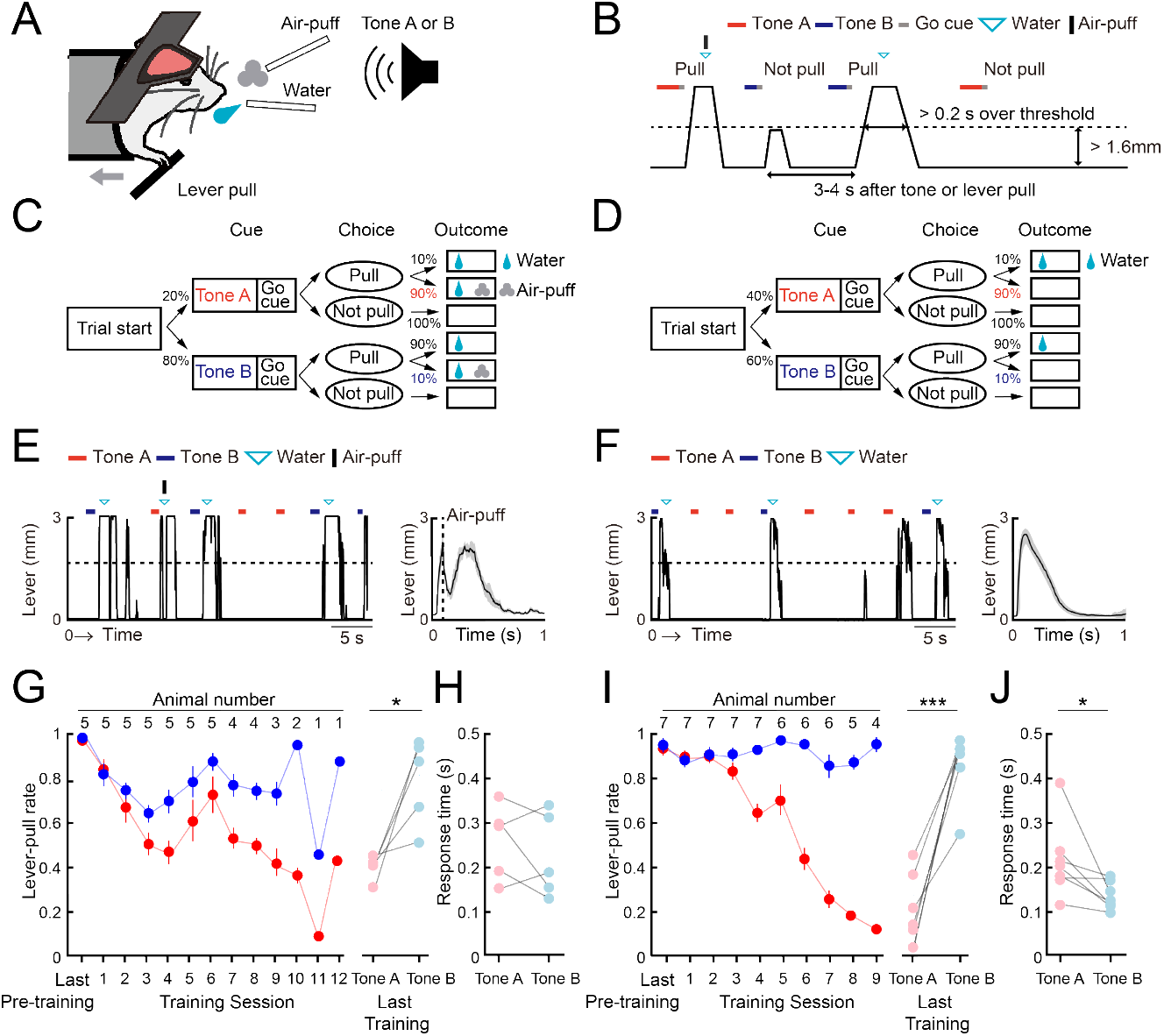
Both positive and negative punishments suppress the reward-seeking behavior of mice. **(A)** Behavioral task setup. **(B)** Schematic diagram of the air-puff task and the omission task. Black line, lever trajectory. Red horizontal bar, tone A presentation. Blue horizontal bar, tone B presentation. Gray horizontal bar, go cue presentation. Cyan inverted-triangle, water reward delivery. Black vertical bar, air-puff delivery. The air-puff was not delivered in the omission task. **(C)** Structure of the air-puff task. The air-puff was delivered at 90% of tone A trials with lever-pull and 10% of tone B trials with lever-pull. The water reward was delivered at 100% of trials with lever-pull regardless of tone type. **(D)** Structure of the omission task. The water reward was omitted at 90% of tone A trials with lever-pull and 10% of tone B trials with lever-pull. **(E)** Left, representative lever trajectories for one mouse (AM1) in the air-puff task in the last (12th) training session. Right, trial-averaged lever trajectories with air-puff in the last training session aligned to the onset of the go cue presentation. Both tone A and B trials for all mice are included. The shading indicates ± standard error of the mean (SEM) across the animals (n = 5). **(F)** Left, representative lever trajectories for one mouse (OM1) in the omission task in the last (8th) training session. Right, trial-averaged lever trajectories with reward omission in the last training session aligned to the onset of the go cue presentation. Both tone A and B trials for all mice are included. The shading indicates ±SEM across the animals (n = 7). **(G, I)** Left, averaged time course of the lever-pull rate in tone A (red) and B (blue) trials in the last pre-training session and during the training sessions in the air-puff task (n = 5) (G) and omission task (n = 7) (I). The number above each session indicates the number of animals in each session. Right, the lever-pull rate in the last training session in tone A (red) and tone B (blue) trials. Light colored dots indicate individual mice. *p = 0.0134, ***p = 0.0008, paired *t* test. **(H, J)** Response time in tone A (red) and B (blue) trials in the last training session of the air-puff task (H) and the omission task (J). Light colored dots indicate individual mice. In (H), the response time was not different between tone A and B trials (p = 0.58, n = 5, paired *t* test), whereas in (J), it was different (*p = 0.035, n = 7).

After pre-training with the reward delivery at 100% probability after lever-pulls in both tone A and B trials without the air-puff, the air-puff was introduced to the air-puff task and the reward omission was introduced to the omission task. In both tasks, as the training progressed, the lever-pull rate (the number of successful lever-pull trials divided by the number of total trials except for those with lever pulling before the go cue presentation) became lower in tone A trials than in tone B trials (Figures 1E and 1F). The air-puff disturbed the lever-pull trajectory (Figure 1E). If mice showed lever-pull rates < 0.5 in tone A trials and > 0.5 in tone B trials for two consecutive training sessions (threshold sessions), we considered that they had learned the task and they were subsequently subjected to two sessions with pharmacological experiments, one with injection of artificial cerebrospinal fluid (ACSF) into the mPFC and one with injection of muscimol into the mPFC (ACSF and muscimol sessions, respectively). The mice that performed all the training, ACSF, and muscimol sessions with the air-puff (n = 5) and omission (n = 7) tasks were used for the analyses in the current study.

In these mice, the lever-pull rate in tone A trials gradually decreased in both air-puff and omission tasks (Figures 1E and 1G), whereas the lever-pull rate in tone B trials with the air-puff task slightly decreased in the first few sessions and then gradually increased (Figure 1G). The lever-pull rate in tone B trials in the omission task remained high (> 0.8) (Figure 1I). As a result, in the last training session, the lever-pull rates between tone A and B trials were different in both air-puff and omission tasks (Figures 1G and 1I). Response time (the latency between cue presentation and the onset of movement) is frequently used to estimate attention and reward expectations^23^, and in the last training session, this time was similar between tone A and B trials for the air-puff task (Figure 1H), but was longer in tone A trials than in tone B trials for the omission task (Figure 1J). This suggests that the response time did not simply reflect the magnitude of punishment expectation, but that it reflected the magnitude of reward expectation to some extent.

### The air-puff is well modeled as positive punishment in the framework of reinforcement learning

To better understand how positive punishment (air-puff) and negative punishment (reward omission) affected the choice behavior (pull or non-pull) of the mouse, we constructed *Q*-learning models with a maximum of five parameters^5,24^ to predict the choice behavior during the training sessions in the air-puff and omission tasks (Table S1; see STAR Methods for details). On the basis of our previous study^5^, we assumed that these tasks included two choices, pull and non-pull, and there were values of pulling the lever (*Q*_pull_) and non-pulling of the lever (*Q*_non-pull_) for both tone A and B trials in each task. The pull-choice probability in each trial was determined from the sigmoidal function of the difference between *Q*_pull_ and *Q*_non-pull_ in that trial. To update *Q*-values, we introduced the following terms: a reward value term (*κ*_r_) that increased *Q*_pull_ after the reward acquisition, a punishment term^25^ (*κ*_p_) that reduced *Q*_pull_ after the punishment was delivered (air-puff stimulus in the air-puff task and reward omission in the omission task), and a saving term (*Ψ*, equal to a covert reward for non-pull) that increased *Q*_non-pull_ after the lever was not pulled. We introduced the term *Ψ* because we previously found that *Ψ* well explained the decrease in the pull probability in tone A trials during training for a similar omission task^5^. We also added a learning rate (*α*_l_) and forgetting rate (*α*_f_), which represent the change rates of *Q*_pull_ and *Q*_non-pul ^26,27^_. In all models, *κ*_r_ and *α*_l_ were included. Seven models were constructed depending on how the use of *κ*_p_, *Ψ*, and *α*_f_ was combined (Table S1). To estimate the predictive performance of the models, we used Akaike’s information criterion (AIC) and Bayesian information criterion (BIC)^28^.

In both tasks, the model with *Ψ* and *α*_f_ and without *κ*_p_ (S-F model) and the model with *κ*_p_, *Ψ*, and *α*_f_ (P-S-F model) appeared to explain the choice behavior with trial-to-trial variability better than the other models (Figures 2A, 2B, and S1A–S1D). In the air-puff task, the P-S-F model was the best-fitting model in five and four out of five mice according to the AIC and BIC scores, respectively (Figure 2C). By contrast, in the omission task, the S-F model was the best-fitting model in six and five out of seven mice according to the respective AIC and BIC scores (Figure 2D). These results indicate that the air-puff effect was well modeled as an overt punishment, with the punishment term *κ*_p_ reducing the value of pull-choice, but the reward omission not doing this.

**Figure 2.**
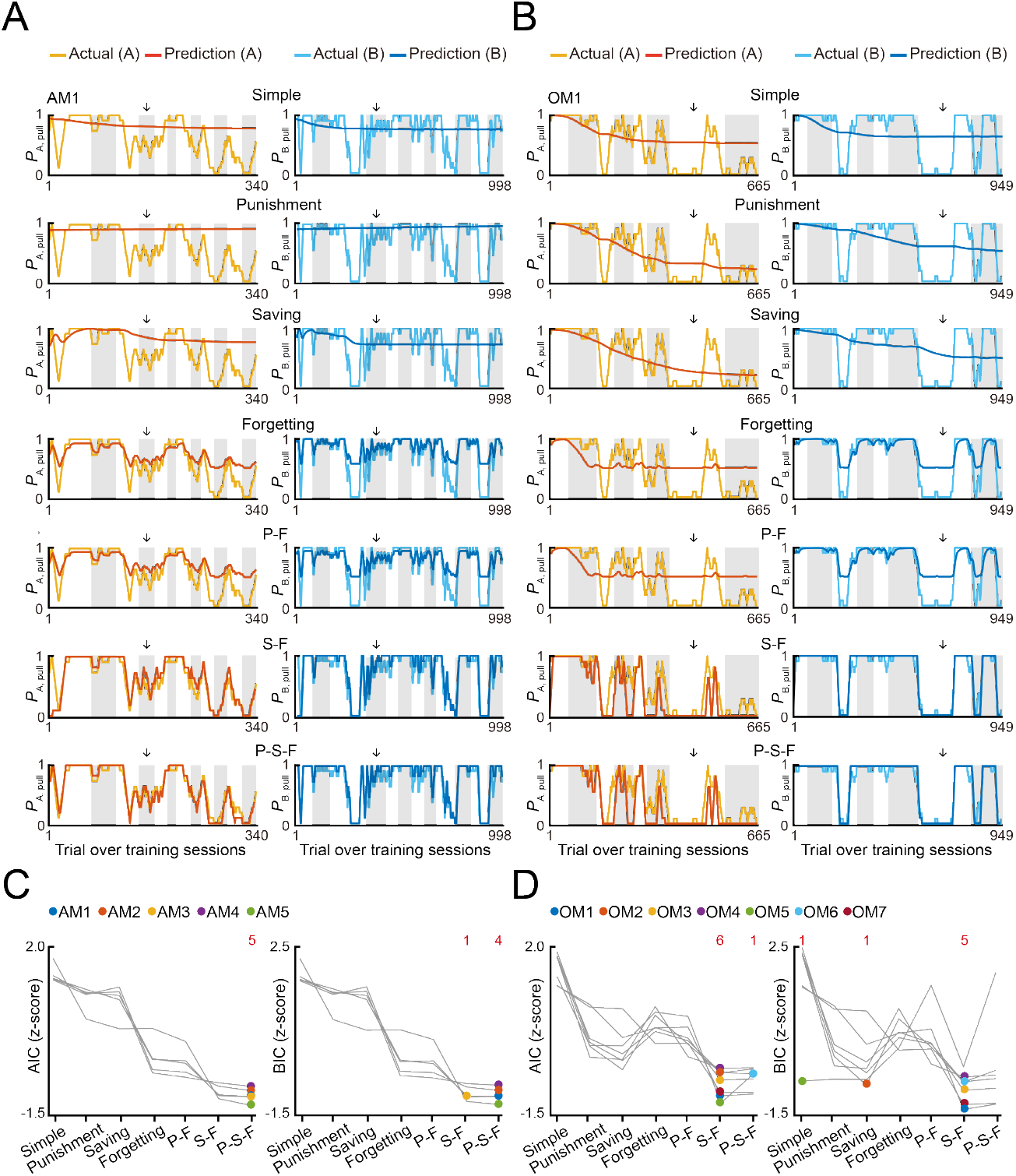
Reinforcement learning models to explain the pull-choice behavior in the air-puff and omission tasks. **(A, B)** Time course of predicted pull-choice probabilities in seven types of reinforcement learning model (simple, punishment, saving, forgetting, P-F, S-F, and P-S-F models) in tone A (left, red) and tone B (right, blue) trials for one mouse (AM1) in the air-puff task (A) and for one mouse (OM1) in the omission task (B). The 10-trial moving averages of the actual pull choice in tone A (yellow) and tone B (cyan) trials are overlaid. Even sessions are shaded. Arrows indicate the second threshold sessions. See also Figure S1 and Table S1. **(C, D)** Z-scored AIC (left) and BIC (right) of seven types of reinforcement learning models in the air-puff task (C) and omission task (D). Colored dots indicate the model with the best score for each mouse. Red numbers above the graph indicate the number of dots for the corresponding model. Each color indicates a different animal (AM1–AM5 in the air-puff task and OM1–OM7 in the omission task).

To examine whether the best-fitting model was also the best for generating pull-choice behavior similar to that shown by the actual mice, we conducted a model simulation^2,5,24,29^. For each mouse, we used the fitted parameters in the S-F and P-S-F models to simulate the lever-pull choice (1, pull; 0, non-pull) in each trial in the order of the actual tone A and B trials (Figures 3A, 3D, and S2A–S2D). In the air-puff task, the simulation with the P-S-F model better regenerated the time course of the actual choice behaviors in both tone A and B trials than that with the S-F model (Figure 3B). The goodness of the generative performance of each model was estimated by calculating the root mean squared error (RMSE) between the actual choice and the simulated choice. The RMSE was smaller with the P-S-F model than with the S-F model (Figure 3C). Thus, the P-S-F model was better than the S-F model as both a fitting model and a generative model. In the omission task, both S-F and P-S-F models well reproduced the choice behaviors (Figure 3E), and there was no significant difference in the RMSE of these two models (Figure 3F). Considering that there were less parameters in the S-F model (4) than in the P-S-F model (5), we concluded that the S-F model explained the choice behavior in the omission task better than the P-S-F model.

**Figure 3.**
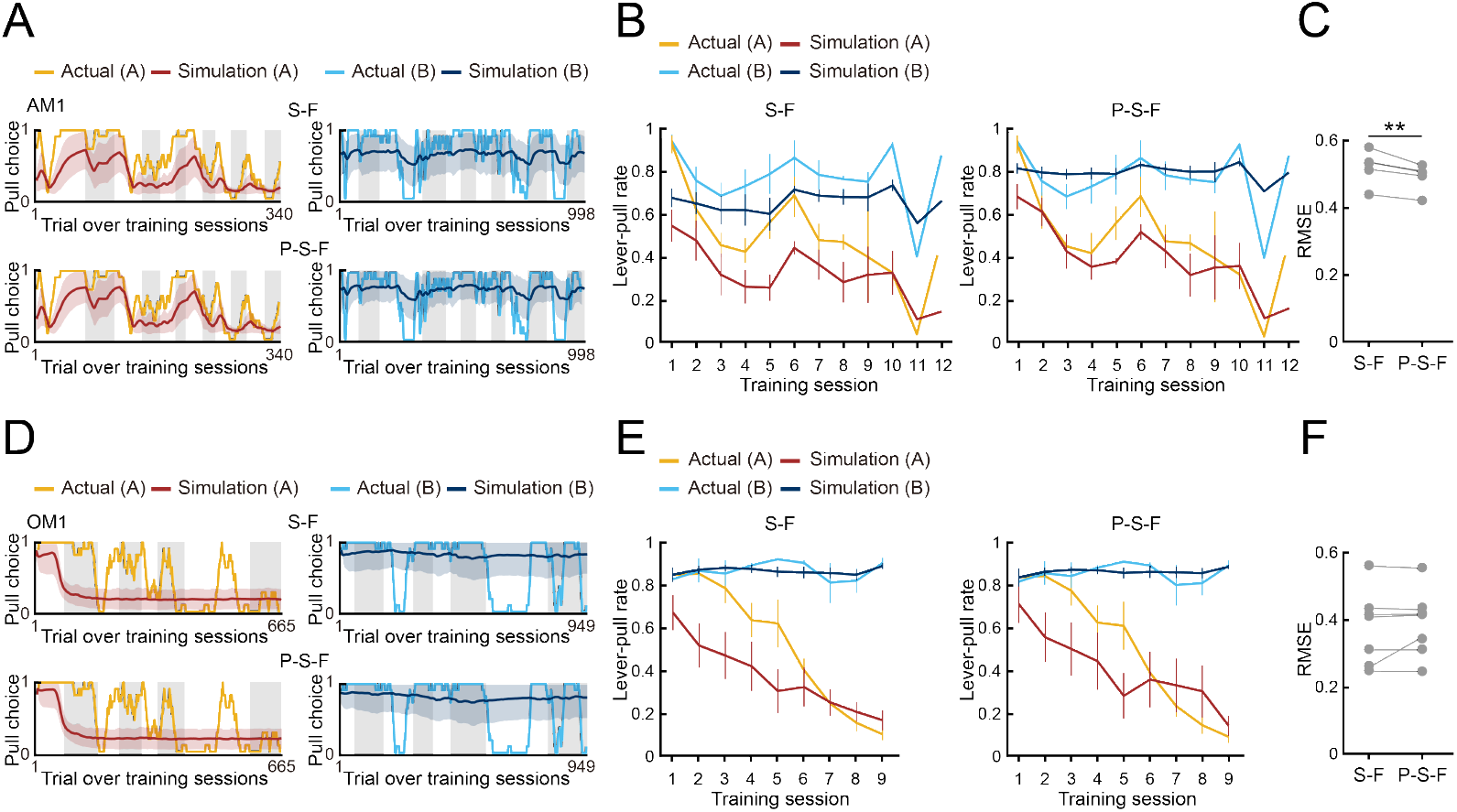
Simulation of choice behavior with S-F and P-S-F models. **(A, D)** Simulated pull-choice behaviors with S-F (upper) and P-S-F (lower) models in tone A (left) and tone B (right) trials for the mouse AM1 in the air-puff task (A) and for the mouse OM1 in the omission task (D). For each mouse, simulation was repeated 1000 times and the choice behavior was averaged in each trial (0–1). The 10-trial moving averages of the simulated pull choice in tone A (dark red) and tone B (dark blue) trials are shown. The shading indicates ±SD across 1000 simulations. The yellow and cyan traces represent the 10-trial moving averages of the actual pull choices in tone A and tone B trials, respectively. Even sessions are shaded.**(B, E)** Mouse-averaged actual (tone A trials, yellow; tone B trials, cyan) and simulated (tone A trials, dark red; tone B trials, dark blue) lever-pull rates across training sessions in the air-puff task (n = 5) (B) and the omission task (n = 7) (E). For each mouse, the simulated lever-pull rate was averaged over 1000 simulations. The error bar indicates ±SEM across the animals. **(C, F)** The RMSE between the simulated and actual pull choices in the air-puff task (C) and the omission task (F). Each gray line represents an individual mouse in S-F and P-S-F models. For each mouse, the RMSE was averaged over 1000 simulations. In (C), **p = 0.0083, paired *t* test (n = 5). In (F), p = 0.3950 (n = 7).

Next, we compared *Q*_pull_ and *Q*_non-pull_ between the P-S-F model in the air-puff task and S-F model in the omission task. *Q*_pull_ in tone A trials (*Q*_A, pull_) decreased and *Q*_pull_ in tone B trials (*Q*_B, pull_) was basically high in both the air-puff and omission tasks throughout the training sessions and *Q*_B, pull_ was higher than *Q*_A, pull_ in the last session in both tasks (Figures 4A–4D). By contrast, *Q*_non-pull_ in tone A trials (*Q*_A, non–pull_) and *Q*_non-pull_ in tone B trials (*Q*_B, non–pull_) in the air-puff and omission tasks fluctuated during the training sessions, and *Q*_A, non–pull_ and *Q*_B, non–pull_ was not different in the last session in both tasks (Figures 4A–4D). When the parameters were compared between the two tasks (Figures 4E–4I), the learning rate (*α*_l_) was higher in the air-puff task than in the omission task, the reward value (*κ*_r_) was lower in the air-puff task than in the omission task, and the saving term (*Ψ*) was lower in the air-puff task than in the omission task, but the difference was not significant.

**Figure 4.**
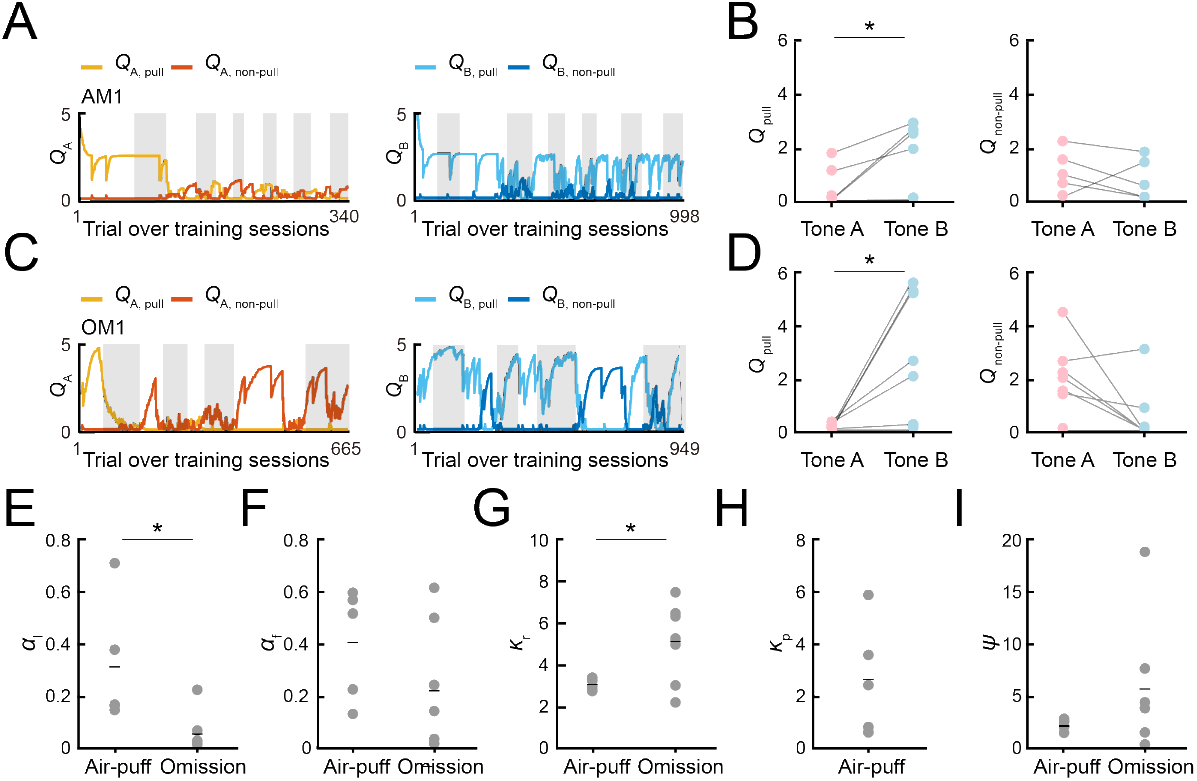
*Q*-values and model parameters in the best-fitting models. **(A, C)** Time courses of *Q*_A, pull_ (yellow) and *Q*_A, non–pull_ (red) (left), and *Q*_B, pull_ (cyan) and *Q*_B, non–pull_ (blue) (right), for the mouse AM1 in the air-puff task (A) and for the mouse OM1 in the omission task (C). **(B, D)** Left, *Q*_A, pull_ and *Q*_B, pull_ averaged within the last training session in the air-puff task (n = 5) (B) and the omission task (n = 7) (D). Right, *Q*_A, non–pull_ and *Q*_B, non–pull_ averaged within the last training session in the air-puff task (B) (n = 5) and the omission task (D) (n = 7). *Q*_A, pull_ vs. *Q*_B, pull_ in the air-puff task, *p = 0.0184; *Q*_A, non–pull_ vs. *Q*_B, non–pull_ in the air-puff task, p = 0.4780; *Q*_A, pull_ vs. *Q*_B, pull_ in the omission task, *p = 0.0195; *Q*_A, non–pull_ vs. *Q*_B, non–pull_ in the omission task, p = 0.0559, paired *t* test. **(E–I)** Model parameters in the P-S-F model for the air-puff task (n = 5) and in the S-F model for the omission task (n = 7). The comparison between the air-puff and omission tasks was conducted with an independent *t* test. (E) Learning rate (*α*_l_). *p = 0.0218. (F) Forgetting rate (*α*_f_). p = 0.2008. (G) Reward value (*κ*_r_). *p = 0.0456. (H) Strength of aversion to the air-puff (*κ*_p_). (I) Non-pull preference (*Ψ*). p = 0.2700. Short horizontal lines indicate animal-averaged values (n = 5 for air-puff task, n = 7 for omission task).

Thus, in the reinforcement learning models, there were differences between the air-puff and omission tasks, not only in respect to whether *κ*_p_ was included or not, but also in the values of *α*_l_ and *κ*_r_. The high *α*_l_ in the air-puff task might reflect the fact that the air-puff increased the attention to the task structure to avoid the positive punishment as soon as possible. The low *κ*_r_ in the air-puff task might reflect the fact that the rarity value of the reward per trial was lower in the air-puff task than in the omission task^5^. It is also possible that the air-puff (positive punishment) suppressed *κ*_r_ more strongly than the reward omission (negative punishment) did. The reward omission was not represented as the overt punishment *κ*_p_, but it might increase the covert reward for non-action *Ψ* more than the positive punishment, and the high *Ψ* would increase the non-pull choice in tone A trials. As a result, *Q*_A, non–pull_ might tend to become larger than *Q*_B, non–pull_ (Figure 4D) as previously suggested^5^. These results indicate that although the time course of the lever-pull rate in tone A and B trials during training sessions was relatively similar between these tasks, the learning process related to the choice behavior might be substantially different.

### mPFC inactivation promotes lever-pull choice in the air-puff task, but not in the omission task

Next, we examined how the mPFC contributed to the choice behavior in the air-puff and omission tasks. ACSF or muscimol was injected into the bilateral mPFC before the start of the ACSF or muscimol session, respectively. The injection was mainly targeted into the prelimbic region (ML 0.2 mm, AP 1.8 mm, DV 1.5 mm; Figures 5A, 5B, S3A, and S3D). The order of the ACSF and muscimol sessions was randomized for each mouse.

**Figure 5.**
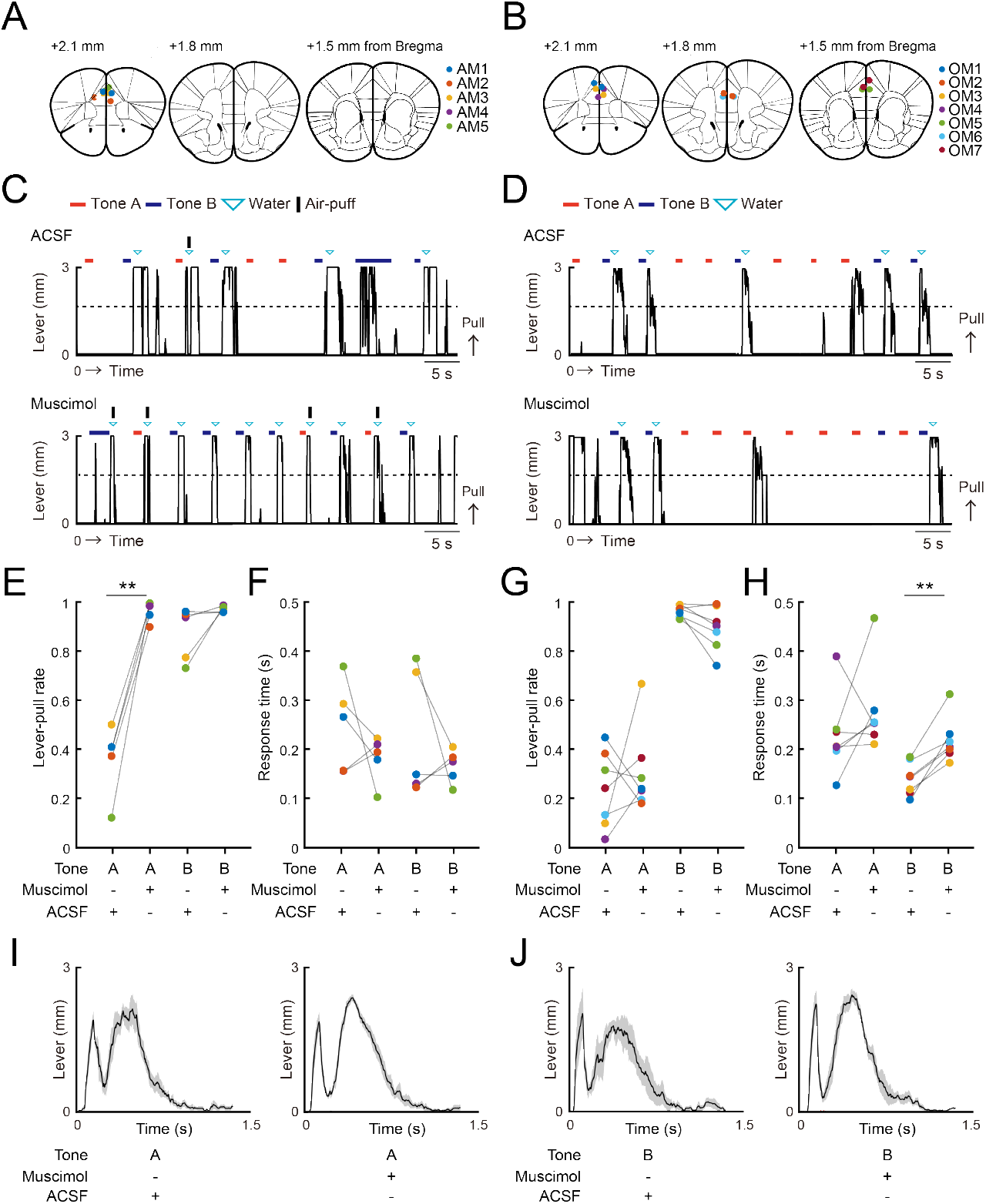
mPFC inactivation promotes lever-pull choice in the air-puff task, but not in the omission task. **(A, B)** The injection sites for ACSF and muscimol in the air-puff task (A) and the omission task (B). The injection sites were inferred from fluorescence labeling in brain slices. Each colored dot represents an individual mouse (AM1–AM5 and OM1–OM7). See also Figure S3. The length from the bregma indicates the distance of the slice along the anterior-posterior axis. The atlas images are taken from The Mouse Brain in Stereotaxic Coordinates^48^. **(C, D)** Representative lever trajectories in the ACSF (upper) and muscimol (lower) sessions for the mouse AM1 in the air-puff task (C) and for the mouse OM1 in the omission task (D). Black line, lever trajectory. Red bar, tone A presentation. Blue bar, tone B presentation. Cyan inverted-triangle, water reward delivery. Black horizontal bar, air-puff delivery. See also Figures S3 and S5. **(E)** Lever-pull rate in tone A (red) and B (blue) trials in the ACSF and muscimol sessions of the air-puff task (n = 5). Each colored line represents an individual mouse (AM1–AM5). ACSF vs. muscimol in tone A trials, **p = 0.0051 by paired *t* test. ACSF vs. muscimol in tone B trials, p = 0.2233 by paired *t* test (n = 5). **(F)** Response time in tone A (red) and B (blue) trials in the ACSF and muscimol sessions of the air-puff task (n = 5). Each colored line represents an individual mouse (AM1–AM5). ACSF vs. muscimol in tone A trials, p = 0.3216 by paired *t* test. ACSF vs. muscimol in tone B trials, p = 0.3745 by paired *t* test (n = 5). **(G)** Lever-pull rate in tone A (red) and B (blue) trials in the ACSF and muscimol sessions of the omission task (n = 7). Each colored line represents an individual mouse (OM1– OM7). ACSF vs. muscimol in tone A trials, p = 0.4917 by paired *t* test. ACSF vs. muscimol in tone B trials, p = 0.1627 by paired *t* test (n = 7). **(H)** Response time in tone A (red) and B (blue) trials in the ACSF and muscimol sessions of the omission task (n = 7). Each colored line represents an individual mouse (OM1–OM7). ACSF vs. muscimol in tone A trials, p = 0.2829 by paired *t* test. ACSF vs. muscimol in tone B trials, **p = 0.0012 by paired *t* test (n = 7). **(I, J)** Lever trajectories averaged across trials and aligned to the onset of the go cue presentation in tone A with the air-puff stimulus (I) and tone B trials with the air-puff stimulus (J) in the ACSF and muscimol sessions of the air-puff task. The shading indicates ±SEM across the animals (n = 5).

The ACSF injection did not appear to change the choice behavior in either task (Figures 5C and 5D). By contrast, the muscimol injection increased the pull choice in tone A trials in the air-puff task (Figures 5C and 5E), although the response times in both tone A and B trials did not show a large change (Figure 5F). However, the muscimol injection did not change the pull choice in tone A trials in the omission task (Figures 5D and 5G). Furthermore, the response time in tone A trials was similar between the muscimol and ACSF sessions, whereas in tone B trials it was longer in the muscimol session than in the ACSF session (Figure 5H).

One possible mechanism to explain the change in the lever-pull rate following muscimol injection in the air-puff task was that mPFC inactivation reduced the sensory response to the air-puff. However, in the muscimol and ACSF sessions, the lever-pull movement was similarly disturbed after the air-puff with both tone types (Figures 5I and 5J). This result suggests that although the mice with mPFC inactivation sensed the air-puff in terms of the withdrawal response, they came to frequently pull the lever regardless of the probability of the air-puff stimulus.

### mPFC inactivation causes hypersensitivity to reward in the air-puff task

Here, we hypothesized that in the air-puff task, the mPFC inactivation changed some internal state that was specifically related to one of the variables in the best-fitting reinforcement learning model, but that this internal state change did not happen in the omission task. In particular, we suspected that the strength of aversion to the air-puff (κ_p_) might be reduced in the muscimol session with the air-puff task, and therefore the value of the lever-pull increased. To test this, we investigated which model parameter best explained the change in choice behavior in the muscimol session. First, we used the parameter set that best explained the choice behavior during the training sessions to simulate the choice behavior in the subsequent ACSF and muscimol sessions, using the P-S-F and S-F models to simulate the choice behavior in the air-puff and omission tasks, respectively. Second, to identify the parameter that best explained the change in the choice behavior in response to the muscimol injection, the value of only one of the parameters (α_l_, α_f_, κ_r_, κ_p_, and Ψ) was changed at a time to minimize the difference between the actual and simulated choice behaviors (Figures 6A–6F). The parameter that minimized the RMSE between the actual and simulated choice behavior was considered to be the injection-related parameter (Figures 6E and 6F). We refer to this second calculation process as parameter optimization.

**Figure 6.**
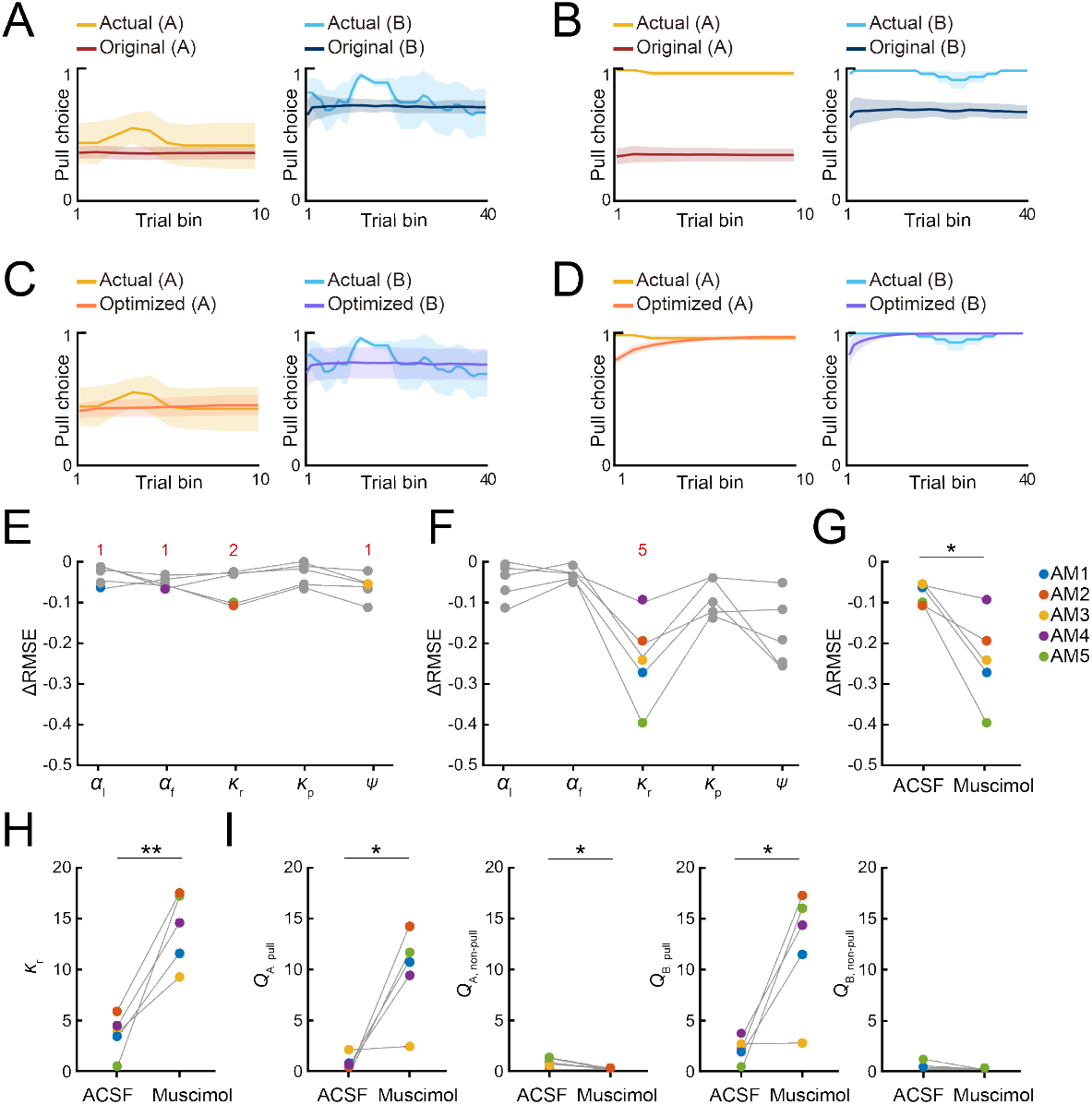
mPFC inactivation causes hypersensitivity to reward in the air-puff task. **(A, B)** Simulated pull-choice behaviors in the P-S-F model with the best parameter set for the training sessions in the ACSF session (A) and muscimol session (B) in the air-puff task. For each mouse, the simulation was repeated 1000 times and the choice behavior was averaged in each trial (0–1). To average the choice behavior across the mice with different trial numbers, the trials in the session were divided into 10 bins in tone A trials and 40 bins in tone B trials for each mouse, and the choice behavior was averaged within each bin. Finally, for each bin, the choice behavior was averaged over the mice. The yellow and cyan traces represent the actual pull choices, whereas the dark red and dark blue traces represent the pull-choice simulated with the original parameters. The shading indicates ±SEM across the animals (n = 5). **(C, D)** Simulated pull-choice behaviors in the P-S-F model after the parameter optimization in the ACSF session (C) and muscimol session (D) in the air-puff task. The yellow and cyan traces represent the actual pull choices, whereas the orange and purple traces represent the pull-choice simulated with the parameters after parameter optimization. The shading indicates ±SEM across the animals (n = 5). **(E, F)** Improvement of the simulation of the choice behavior after the parameter optimization of each parameter in the ACSF session (E) and muscimol session (F) in the air-puff task. Each line indicates a different animal (AM1–AM5). Each colored dot indicates the parameter that minimized the error between the actual and simulated choice behaviors (ΔRMSE) for each mouse. Red numbers indicate the number of the colored dots in the corresponding parameter. **(G)** The change in the RMSE (ΔRMSE) resulting from parameter optimization in the simulation of the behaviors in the ACSF and muscimol sessions. Each colored line indicates an individual mouse. *p = 0.0237, paired *t* test. **(H)** κ_r_ after the parameter optimization in the ACSF and muscimol sessions. Each color indicates a different animal (AM1–AM5). **p = 0.0052, paired *t* test. **(I)** *Q*_A, pull_, *Q*_A, non–pull_, *Q*_B, pull_, and *Q*_B, non–pull_ estimated with the parameters obtained after parameter optimization in the ACSF and muscimol sessions. Each line represents an individual mouse. Each dot indicates the average within each session. *Q*_A, pull_, *p = 0.0185; *Q*_A, non–pull_, *p = 0.0304; *Q*_B, pull_, *p = 0.0214; *Q*_B, non–pull_, p = 0.1291, paired *t* test (n = 5). Each colored line indicates an individual mouse (AM1–AM5).

In the air-puff task, the parameter set that best explained the behaviors during the training sessions well simulated the choice behavior in the ACSF session (Figures 6A and S3B), and the parameter optimization did not greatly improve the accuracy of the simulation (Figures 6C, 6E, and S3B). The injection-related parameters were different across the five animals (Figure 6E). These results indicate that the behavior in the ACSF session largely inherited the internal states present during the training session. By contrast, the parameter set for the training sessions did not well simulate the behaviors in the muscimol session (Figures 6B and S3C). However, after the parameter optimization, the increased lever-pull choice in tone A trials was well simulated (Figures 6D and S3C). In four of five mice, the goodness of the simulation was substantially improved (Figure 6F). The change in the RMSE (ΔRMSE) resulting from the parameter optimization was more pronounced in the muscimol session than in the ACSF session (Figure 6G). In this optimization, the injection-related parameter was the same across all five mice: the reward value *κ*_r_ increased substantially in all mice (Figure 6H). As a result, the action values (*Q*_pull_) in both tone A and B trials increased (Figure 6I).

The difference between the actual and simulated choice behaviors with the parameter set for the training sessions was particularly apparent in the early part of the muscimol session (Figures 6B, 6D, and S3C). Thus, the increase in *κ*_r_ might just be sufficient to minimize this difference as rapidly as possible. To test this hypothesis, we also considered the potential effects of muscimol injection on the initial *Q*-values. Assuming that these values reached a steady-state value after the training sessions, we calculated them from the parameter set (see Methods for details). In this case, the decrease in *κ*_p_ and the decrease in *Ψ*, as well as the increase in *κ*_r_, should increase the action value and pull-choice probability, even in the first tone A trial. However, even if we included the initial action value in the parameter optimization, *κ*_r_ was chosen as the injection-related parameter in all five mice (Figures S4A–S4C). Regardless of whether or not the initial action values were considered in the parameter optimization, *κ*_r_, *Q*_A,pull_, and *Q*_B,pull_ were significantly larger in the muscimol session than in the ACSF session, whereas *Q*_A,non-pull_ was significantly smaller in the muscimol session (Figures 6H, 6I, S4D, and S4E). In summary, for the air-puff task, the parameters that best simulated the choice behavior in the ACSF sessions varied across the mice, whereas the simulation of the choice behavior in the muscimol session was most improved by increasing the reward value (*κ*_r_) for all mice. The model therefore suggests that the increase in *κ*_r_ increased the action value (*Q*_pull_), so that the pull-choice probability was high from the start, even though the probability of the air-puff stimulus was as constantly high in tone A trials as in the training sessions.

In comparison, the injection-related parameters for the muscimol and ACSF sessions in the omission task varied across the mice (Figures S3D–S3F and S5A–S5F). ΔRMSE was also small after the parameter optimization (Figures S5E–S5G). This suggests that the internal state related to the S-F model parameters in the omission task was not changed by mPFC inactivation. This is consistent with the result that mPFC inactivation did not affect the choice behavior in the omission task.

## Discussion

In the present study, we examined how the reward-seeking behavior of mice differed under the risks of positive (air-puff stimulus) and negative (reward omission) punishments. Both types of punishment weakened the lever-pull action of mice in the trials with a higher probability of air-puff stimulus and reward omission. The positive punishment (air-puff) was well modeled as the overt punishment in the framework of reinforcement learning, from both predictive and generative aspects. In addition, inactivation of mPFC increased the pull action in the trials with a higher probability of the air-puff stimulus. However, the mPFC inactivation had no effect on the low pull-choice in response to the tone with a high probability of reward omission (equal to a low probability of reward delivery). Furthermore, we developed an analytical method to change one of the parameters of the reinforcement learning to explain the change in choice behavior under the acute areal inactivation. This method suggested that the mPFC inactivation increased the reward value under risk of positive punishment.

The necessity of the mPFC for decision-making under risk of positive punishment is well established^9-18^. For example, Friedman et al. ^15^ showed that inhibition of neurons in the prelimbic region increases the probability of rats choosing a high-risk option (rats were exposed to brighter light but at the same time they could get sweeter milk chocolate) only when there are cost-benefit conflicts, and prelimbic inhibition does not affect the behavior when there is no conflict. Our result for the behavioral change in the muscimol session with the air-puff is consistent with these previous studies. In contrast to the air-puff task, mPFC inactivation did not change the choice behavior in the omission task. This result is consistent with another study that found that mPFC inactivation did not change the choice between a certain small reward and a risky large reward^19^. These results suggest that similar mPFC activity occurs in decision-making tasks of whether or not to act, as well as in many tasks involving the choosing of one action from multiple actions. However, there is currently no unanimous answer as to which parameter in the reinforcement learning model best explains the effect of the mPFC inactivation in both task structures. Here, we explained the change in the choice behavior in terms of a change in a reinforcement learning model parameter. Previous studies applied such an approach to primate behavior^29-33^, but the current study applied it to rodent behavior. In addition, no specific change in the sensitivity to reward (or subjective value of the reward) has been reported in areal inactivation experiments in the model animal. Our analytical method can be applied to many other types of decision-making tasks to reveal the parameter that is represented by each brain region, or even each cell type, in the reinforcement learning model for the decision-making process.

Impairment of the mPFC is frequently related to drug addiction^34-39^, but the reinforcement learning model has not produced consistent results for explaining human decision-making behaviors. For example, cocaine users show a decreased learning rate^37^, heroin users show reduced loss aversion, amphetamine users show increased sensitivity to reward^38^, and cannabis users show increased sensitivity to gains and decreased sensitivity to losses^39^. In these drug users, long-lasting effects of the drugs would impair many aspects of their brain circuits, so that consistent results may not be obtained. By contrast, our study on acute mPFC inactivation clearly suggests that the increased sensitivity to reward value, but not the decreased aversion strength to punishment, may be the most relevant to the mPFC-impaired behavior. This result is consistent with observations in patients with VM lesions, who show hypersensitivity to reward^10^. However, since we changed only one parameter value, we do not exclude the possibility that parameters other than the reward value would also be affected. Alternatively, mPFC inactivation might change the animal’s behavioral strategy from the decision making-based behavior to a habitual behavior that cannot be explained by the reinforcement learning framework. However, in the omission task, the pull choice in tone A trials remained probabilistic, as in the other sessions, and the reaction time in tone B trials was rather lengthened. Thus, the behavioral change in the muscimol session of the air-puff task did not simply reflect the habitual behavior; the behavioral change needed the positive punishment. To directly test whether the increase in the pull choice was addictive and habitual, one could examine whether the lever-pull choice continues or not after the lever-pull is devaluated in the middle of the muscimol session. The rodent mPFC consists of prelimbic and infralimbic regions and the ACC. These regions have distinct functions in reward-seeking behaviors under the risk of punishment^9,15,40^. Considering that dmPFC and vmPFC in humans signal risk and outcome values, respectively^9^, the behavioral change by the inactivation of mPFC in the air-puff task might be caused by the inactivation of both dorsal and ventral parts of mPFC, although the injection site was targeted to the prelimbic region. It is necessary to clarify how each region is related to the choice behavior and reward sensitivity under risk of positive punishment.

If mPFC reduces the sensitivity to reward under risk of positive punishment, what is the mechanism? mPFC and the ventral tegmental area (VTA) are strongly related to positive punishment on different timescales^1^. VTA dopaminergic neurons display phasic excitatory responses tightly linked to each task event (cue, action, punishment, and reward), and might be involved in the short-timescale real-time neural processing of punishment to promote rapid behavioral adaptation. By contrast, mPFC neurons show the punishment-related responses that are longer than the transient activity of VTA dopaminergic neurons, and baseline activity of many of mPFC neurons are modulated^1^. mPFC neurons that project to the dorsomedial striatum maintained the action value over a long timescale in a dynamic foraging task^41^. Thus, mPFC might be responsible for longer-lasting impacts of punishment on motivational and emotional states. Such longer-lasting effects may consistently affect actions over the session, regardless of the presentation of punishment. In this case, the inactivation of mPFC may be better represented in the model as an increase in *κ*_r_ rather than as a decrease in *κ*_p_. From the initial tone A trial in the muscimol session of the air-puff task, the pull was consistently chosen. These results suggest that mPFC maintained the subjective value of the reward within an appropriate range, rather than the aversion strength to the punishment. This might result in the lower *κ*_r_ observed in the air-puff task compared with the omission task (Figure 4G). Alternatively, the low *κ*_r_ in the air-puff task might simply reflect the rarity value of the reward per trial^5^. To test whether mPFC encodes the subjective value of the reward, it is necessary to measure the activity of mPFC in the air-puff task^20,21^ and clarify how its activity and the estimated reward value change when the reward delivery probabilities in tone A and B trials vary.

The activity of the mPFC under risk of positive punishment should be transmitted to other areas. Considering that modulation of the connections from the mPFC to the basolateral amygdala or VTA has a weak effect on reward-seeking behavior^14,15^, its major downstream site would be the striatum, which is critical for updating the action value^42^. Inhibition of glutamatergic projections from the mPFC to the nucleus accumbens core and shell in the rat causes it to choose an option with high benefits-high costs, rather than a choice with low benefits-low costs^15^. The mPFC may suppress the activity of striatal medium spiny neurons by activating striatal interneurons. Medium spiny neurons in the dorsal striatum are largely suppressed under the cost-benefit conflict, with the activity peaks of striatal interneurons preceding the inhibition of peaks of medium spiny neurons^15^. Striatal medium spiny neurons are known to represent reward signals and promote reward-seeking behavior^43^. Therefore, activation of striatal interneurons by the striatum-projecting mPFC neurons may subsequently suppress the striatal medium spiny neurons and prevent the sensitivity to reward from jumping up. If the VTA signal represents the aversion strength to the punishment (*κ*_p_), the signals from mPFC and VTA (*κ*_r_ and *κ*_p_, respectively) may be incorporated when the action value is updated in the striatum. Striatal neurons expressing dopamine D2 receptors are well known to mediate negative feedback through both positive and negative punishment^44–47^. After learning of the risk of negative punishment, these striatal neurons may determine whether to act or not without using the signal from mPFC. However, in the omission task in the present study, it is unknown how the striatal neurons expressing D1 and D2 receptors are related to the choice behavior. To clarify this, optogenetic manipulation and measurement of the striatal neurons and dopaminergic neurons is necessary.

### Limitations of the study

In the current study, we fixed the combination of the reward and air-puff delivery probabilities within and across sessions. It was reported that mPFC inactivation changes choice behavior when the reward omission probability changes within a session^4,19,40^. Thus, mPFC would be required for behavioral flexibility under risk of negative punishment. It is necessary to verify this by varying the reward probability within the session in the current omission task. The probabilities of the tone A and B presentation were also fixed, and were different between the air-puff task and the omission task. It is possible that these fixed conditions created some bias in the choice behavior. In addition, since mPFC has various functions for each sub-region and projection, further examination with a more specific method for modulation, such as chemogenetics and optogenetics, is required.

## Materials and Methods

### Animals

All animal experiments were approved by the Animal Experimental Committee of the University of Tokyo. C57BL/6 mice (male, aged 2–3 months at the start of behavioral training, SLC, Shizuoka, Japan, RRID: MGI:5488963) were used for the experiments. These mice had not been used in other experiments before this study. All mice were provided with food and water ad libitum and housed in a 12:12 hour light-dark cycle (light cycle; 8 AM–8 PM). All behavioral sessions were conducted during the light period.

### Head plate implantation

Mice were anesthetized by intramuscular injection of ketamine (74 mg/kg) and xylazine (10 mg/kg), with atropine (0.5 mg/kg) injected to reduce bronchial secretion and improve breathing, an eye ointment (Tarivid; 0.3% w/v ofloxacin; Santen Pharmaceutical, Osaka, Japan) applied to prevent eye-drying, and lidocaine jelly applied to the scalp to reduce pain. Body temperature was maintained at 36°C–37°C with a heating pad. After the exposed skull was cleaned, an incision was made in the skin covering the neocortex, and a custom head plate (Tsukasa Giken, Shizuoka, Japan) was attached to the skull using dental cement (Fuji lute BC; GC, Tokyo, Japan; and Estecem II; Tokuyama Dental, Tokyo, Japan). The surface of the intact skull was coated with dental adhesive resin cement (Super bond; Sun Medical, Shiga, Japan) to prevent drying. A single intraperitoneal injection of the anti-inflammatory analgesic carprofen (5 mg/kg, Rimadile; Zoetis, NJ, USA) was given after all surgical procedures. Mice were allowed to recover for 3–5 days before behavioral training.

### Behavioral training

After recovery from the head plate implantation, the mice were water-deprived in their home cages. Each mouse received about 1 mL of water per session per day, but they were sometimes given additional water to maintain their body weight at 85% of their initial weight throughout the experiments. The mice were usually trained for five consecutive days per week and were given 1 mL of water on days without training. The behavioral apparatus (sound attenuation chamber, head-fixing frame, body holder, sound presentation system, water-supply system, and integrated lever device) was manufactured by O’Hara & Co., Ltd. (Tokyo, Japan). The lever position was monitored by a rotary encoder (MES-12-2000P), which continuously recorded results at an acquisition rate of 1000 Hz using a NI-DAQ (PCIe-6321; National Instruments, Austin, TX, United States). The sound, water, and air-puff stimulus were controlled using a program written in LabVIEW (2018, National Instruments, RRID: SCR_014325).

### Pre-training

Training was conducted once a day, with the mice in the chamber with a head plate during the training. On the first 2–3 days, a go cue (pink noise, 0.3 s) was presented and a water reward was delivered if the mouse licked a spout within 1 s of the go cue presentation. The mice gradually learned to obtain the water reward by licking the spout after the go cue. They were then moved on to the next task, in which they had to pull the lever more than 1.6 mm for longer than 0.2 s to obtain the reward, rather than just licking the spout. The weight of the lever was fixed at 0.07 N. Over 2–3 days, the mice learned to pull the lever for a duration of more than 0.2 s within 1 s after the go cue was presented. During the last 3–4 days, two tones (6 and 10 kHz pure tones, each of duration 0.8–1.2 s) were alternately presented before the go cue at an equal probability (50%). The next trial started 3–4 s after the last timepoint at which the lever was returned to the home position (after the lever went below the 1.6 mm threshold), or after the presentation of the previous tone cue when the lever did not exceed the threshold. If a mouse pulled the lever during tone presentation, it was counted as an early pull and the tone was prolonged until the mouse stopped pulling the lever and waited for 1.5–2.5 s without pulling the lever again. Pre-training was considered complete when the lever-pull rate was > 90% and the early pull rate was < 30% for both tones.

### Air-puff and omission tasks

In the two-tone lever pull task, either of the tone cues used in the pre-training sessions was randomly presented. The probabilities of tone A and tone B presentations were 20% and 80%, respectively, for the air-puff task, and 40% and 60%, respectively, for the omission task. The mouse heads were fixed in a way that allowed them to pull the lever within 1 s after the cue presentation, as in the pre-training sessions. In the air-puff task, when mice pulled the lever over the threshold (1.6 mm) for longer than 0.2 s, they always received a 4 μL drop in response to either tone cue, and simultaneously received an air-puff (0.3–0.4 Mpa for 20 ms) at probabilities of 90% and 10% in response to tone A and B trials, respectively. Air-puffs were delivered from a needle tip that was located 3–5 mm away from the left eye. In the omission task, mice that pulled the lever in response to tone A and tone B received a 4 μL drop of water at probabilities of 10% and 90%, respectively, but never received the air-puff punishment. Mice that did not pull the lever did not receive water or an air-puff. The next trial started 3–4 s after the last timepoint at which the lever was returned to the home position (after the lever went below the threshold), or after the presentation of the previous tone cue when the lever did not exceed the threshold (1.6 mm). In both tasks, training was considered successful when the lever-pull rates in response to tone A and tone B were < 50% and > 50%, respectively, for two consecutive sessions (threshold sessions) by the tenth (air-puff task) or eighth (omission task) training session. The force required to pull the lever varied from 0.05–0.07 N per mouse, but tended to be greater for the omission task. The early pull rates in the last training session were 0.0 for both tones in the air-puff task (n = 5), and 0.027 ± 0.002 and 0.053 ± 0.006 for trials with tone A and tone B, respectively, in the omission task (n = 7). Trials with early pulls were not included in the behavioral analysis and computational modeling.

### Pharmacological inactivation

Four mice (two in the air-puff task and two in the omission task) performed the sessions with pharmacological inactivation (ACSF and muscimol sessions) without any training session after the second threshold session. The other mice continued to perform 1–8 training sessions until the ACSF and muscimol sessions started. The mice underwent bilateral craniotomies over the prefrontal cortex (ML 0.2 mm, AP 1.8 mm, diameter 2 mm) at least 3 days before the day of injection and the craniotomies were covered with silicone elastomer (Kwik-Cast, World Precision Instruments, FL, USA). Before injection, a glass pipette (3-000-203-G/X, Drummond Scientific Company, PA, USA) was pulled, cut until its outer diameter was around 40 μm, and then backfilled with mineral oil (Nacalai Tesque, Kyoto, Japan). The pipette was fitted to a Nanoject III Programmable Nanoliter Injector (Drummond Scientific Company), and ACSF or muscimol (60 nL; 5 μg/μL; M1523, Sigma-Aldrich, MO, USA) diluted in ACSF was front-loaded. During the intracranial injection, the mice were anesthetized with isoflurane (0.7– 1.4%) inhalation and fixed with a head plate. The pipette tip was inserted into the brain at a depth of 1.5 mm from the cortical surface, and ACSF or muscimol was injected via a Nanoject III at a rate of 8–10 nL/minute. The pipette was maintained in place for 5 minutes after the injection and then slowly withdrawn. Injection into the second hemisphere was always completed within 15 minutes from the beginning of injection into the first hemisphere. The craniotomies were covered with silicone elastomer. Both the air-puff task and the omission task were started 40 minutes after the injection. The order of ACSF and muscimol sessions was randomized for each mouse. After the completion of both sessions, NeuroTrace CM-Dil tissue-labeling paste (N22883, Thermo Fisher Scientific, MA, USA) was injected at the same stereotaxic site to confirm the ACSF and muscimol injection site.

### Histology

Mice were anesthetized by intraperitoneal injection of a mixture of ketamine and xylazine, and were then perfused transcardially with PBS followed by a solution of 4% paraformaldehyde (09154-85, Nacalai Tesque). The brains were removed and stored in the fixative overnight, and were kept at 4°C until being coronally sectioned (100 μm sections) with a Vibratome (VT1000S, Leica, Germany). Sections were mounted in Vectashield mounting medium containing DAPI (H1500, Vector Laboratories, CA, USA) and were imaged with a camera (RETIGA2000, *Q*IMAGING, AZ, USA) attached to a fluorescence microscope (BX53, Olympus, Tokyo, Japan).

### Analysis of behavioral data

The data were analyzed using MATLAB (R2023b; MathWorks, Natick, MA, USA, RRID: SCR_001622). No apparent abnormal behavior was observed on the day after a break (e.g., on a Monday). Therefore, the behavior of the mice was analyzed throughout the first to the last session. To remove the possible period within each session in which mice were less motivated, the behavioral data were determined only from the first trial to the last trial prior to obtaining 60% of the total planned reward^5^. The lever-pull rate was defined as the number of successful lever-pull trials divided by the total number of the trials except for those with early pulls. To visualize the change in pulling or not pulling within each trial, the 10-trial moving-average of the pull-choice (pull was 1 and non-pull was 0) was used.

### Reinforcement learning models

All behavioral data were summarized as binary data with action (to pull or not), cue type, water reward, and air-puff punishment. The trial sequence in each session was determined using the same criterion as the behavioral analyses. The sequences from a single animal were concatenated through all sessions, and then the data series was separated into two sequences consisting of tone A and tone B trials, which were used to model the learning process of the mice.

The model was based on that used in our previous study^5^. The values for pulling and not pulling the lever during the *t*-th trial of each tone cue (*x* ∈ {A, B}) were defined as *Q*_*x*,pull(*t*)_ and *Q*_*x*,non-pull(*t*)_, respectively. *Q*_*x*,pull(*t*)_ was updated according to equations [1] and [3], and *Q*_*x*,non-pull(*t*)_ was updated according to equations [2] and [4].

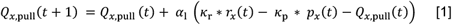

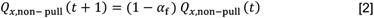

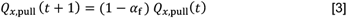

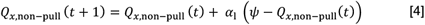

where *α*_l_ (0 < *α*_l_ < 1) was the learning rate, *α*_f_ (0 < *α*_f_ < 1) was the forgetting rate, *κ*_r_ (≥ 0) was the subjective goodness of a water reward (reward value), *κ*_p_ (≥ 0) was the subjective strength of aversion to air-puff punishment, and Ψ (≥ 0) was the goodness of the covert reward, which is assumed to be constantly obtained as a result of a non-pull (i.e., the saving of the cost accompanying the lever-pull)^5,49,50^. In both the air-puff task and the omission task, *r*_*x*_(*t*) corresponded to the water reward (presence, 1; absence, 0) in the *t*-th tone *x* trial. In the air-puff task, *p*_*x*_(*t*) was defined as 1 when mice pulled the lever and an air-puff was delivered in the *t*-th tone *x* trial, whereas *p*_*x*_(*t*) was 0 when mice did not pull the lever or when they pulled the lever but an air-puff was not delivered. In the omission task, *p*_*x*_(*t*) was defined as 1 when the mice pulled the lever but water was not delivered in the *t*-th tone *x* trial, whereas *p*_*x*_(*t*) was 0 when the mice did not pull the lever or when they pulled the lever and water was delivered. The pull-choice probability for the (*t*+1)-th trial for tone *X, P*_*x*,pull_(*t* + 1), was calculated according to equation [5].

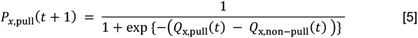

The parameters used for each model are summarized in Table S1.

**Table S1.**
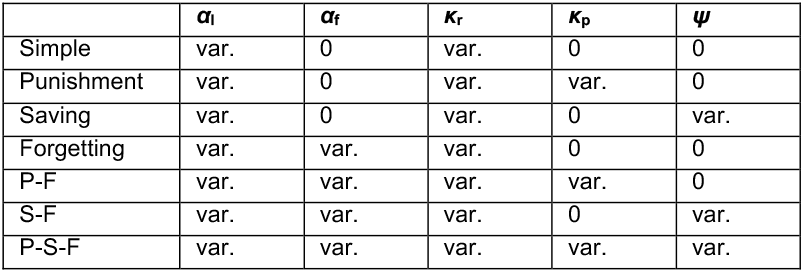
Summary of the free parameters used in each *Q*-learning model, related to Figure 2. Var.: variable.

Maximum log likelihood estimation was used to fit the parameters used in all models. The likelihood (*L*) was calculated according to equation [6]:

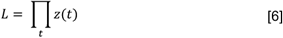

where z(*t*) could be calculated as *P*(*t*) if the lever was pulled, and 1 − *P*(*t*) if the lever was not pulled. The logarithm of this likelihood was multiplied by –1 to allow the use of the *fmincon* function in MATLAB with appropriate lower and upper bounds for each free parameter. For each model, the parameter fitting was repeated 5000 times and the parameters for the fitting that showed the maximum likelihood (*L*_max_) was chosen. To compare the models, Akaike’s information criterion (AIC) and Bayesian information criterion (BIC) were calculated using the following formulas:

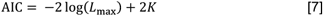

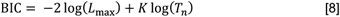

where *K* is the number of free parameters to fit, and *T*_*0*_ is the number of trials used for fitting.

To simulate the choice behavior in the model with the fitted values of the free parameters described above, the same sequences of tones across sessions were used as in the actual settings for each mouse, and the lever-pull choice (pull or non-pull) in each trial was calculated according to the pull-choice probability estimated by equation [5]. When the lever was pulled in the simulated *t*-th trial and was actually pulled in the *t*-th real trial, the actual values of *r*_*x*_(*t*) and *p*_*x*_(*t*) were used. When the lever was pulled in the simulated *t*-th trial but was not actually pulled in the real *t*-th trial, *r*_*x*_(*t*) and *p*_*x*_(*t*) were defined according to the determined probability (the air-puff task, 100% in both tone A and B trials for reward, 90% and 10% in tone A and B trials, respectively, for punishment; the omission task, 10% and 90% in tone A and B trials, respectively, for reward, 0% in both tone A and B trials for punishment). For the simulation of the training processes, the initial values of *Q*_*x*,pull_(1) and *Q*_*x*,non-pull_(1) were the same as those for the fitting. The simulation was repeated 1000 times and the pull choice was averaged in each trial. The goodness of the generative performance of each model was estimated by calculating the root mean squared error (RMSE) between the actual choice and the simulated choice averaged across 1000 simulations. The RMSE was calculated according to equation [9]:

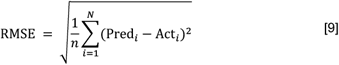

where *N* is the total number of trials including both tone A and tone B trials, Act_*i*_ indicates the actual lever-pull choice (1, pull; 0, non-pull) in the *i*-th trial, and pred_*i*_ indicates the simulated lever-pull choice (1, pull; 0, non-pull) in the *i*-th trial.

where *N* is the total number of trials including both tone A and tone B trials, Act_*i*_ indicates the actual lever-pull choice (1, pull; 0, non-pull) in the *i*-th trial, and pred_*i*_ indicates the simulated lever-pull choice (1, pull; 0, non-pull) in the *i*-th trial.

For visual presentation of mouse-averaged pull-choice behaviors (Figures 6A–6D, S4A, and S5A–S5D), the total number of trials for each mouse was divided into bins (10 in tone A trials and 40 in tone B trials for the air-puff task, 20 in tone A trials and 30 in tone B trials for the omission task), according to the probabilities of tone A and tone B presentations. Then, the average per bin was calculated for each mouse.

To better simulate the pull-choice behavior in the ACSF and muscimol sessions, the parameters in the S-F and the P-S-F models were modulated (parameter optimization). Search ranges for *α*_l_ and *α*_f_ were from 0 to 1 in increments of 0.01, whereas the search ranges for *κ*_r_, *κ*_p_, and *Ψ* were from 0 to 20 in increments of 0.1. One parameter was modulated and the other parameters were fixed at their original fitted values. The last *Q*-values of the model fitting in the training sessions were used as the initial *Q*-values in the ACSF and muscimol sessions. The simulation for each value was repeated 1000 times and the RMSE was calculated according to equation [9]. We considered the parameter that minimized RMSE to be the one that most reflected the effect of the injection.

In the modified version of the parameter optimization in the air-puff task, we assumed that the initial *Q*-values already reflected the value of the parameter that was newly modulated because the injection was conducted before the ACSF or muscimol session started. If equations [1] and [4] were in a steady-state and the mice recognized the expected values of *r*_A_(*t*), *r*_B_(*t*), *p*_A_(*t*), and *p*_B_(*t*) in the air-puff task as the constant values R_A_= 1, R_B_ = 1, p_A_ = 0.9, and p_B_ = 0.1, respectively, the initial *Q*-values should be as follows:

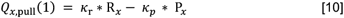

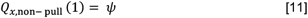

We included these *Q*-values in the simulation and searched for the parameter that minimized the RMSE.

### Quantification and statistical analysis

Groups were compared using independent *t* tests (for Figures 4E–4I) or paired *t* tests (for Figures 1G–1J, 3C, 3F, 4B, 4D, 5E–5H, 6G–6I, S4C–S4E, and S5G), as appropriate. All statistical tests were two-tailed.

## Author Contributions and Notes

M.N., M.K., and M.M. conceptualized the study; M.N., E.Y., and M.K. designed the experiments; M.N. performed the experiments, analyzed the data, and prepared the figures; M.N., M.K., and M.M. interpreted the results; M.N., M.K., and M.M wrote the manuscript; M.M. supervised M.N.

The authors declare no conflict of interest.

## Acknowledgments

The authors thank M. Nishiyama for animal care and breeding, S. Tanimoto for helpful advice on model fitting and simulations, and S. Palminteri for helpful discussion. This work was supported by Grants-in-Aid for Scientific Research on Innovative Areas (17H06309 to M.M.), for Transformative Research Areas (A) (22H05160 to M.M.), and for Scientific Research (A) (19H01037 and 23H00388 to M.M.) from the Ministry of Education, Culture, Sports, Science, and Technology, Japan; and AMED (JP18dm0207027 to M.M.).

