## Supplementary Material for "Medial prefrontal cortex suppresses reward-seeking behavior with risk of positive punishment by reducing sensitivity to reward"

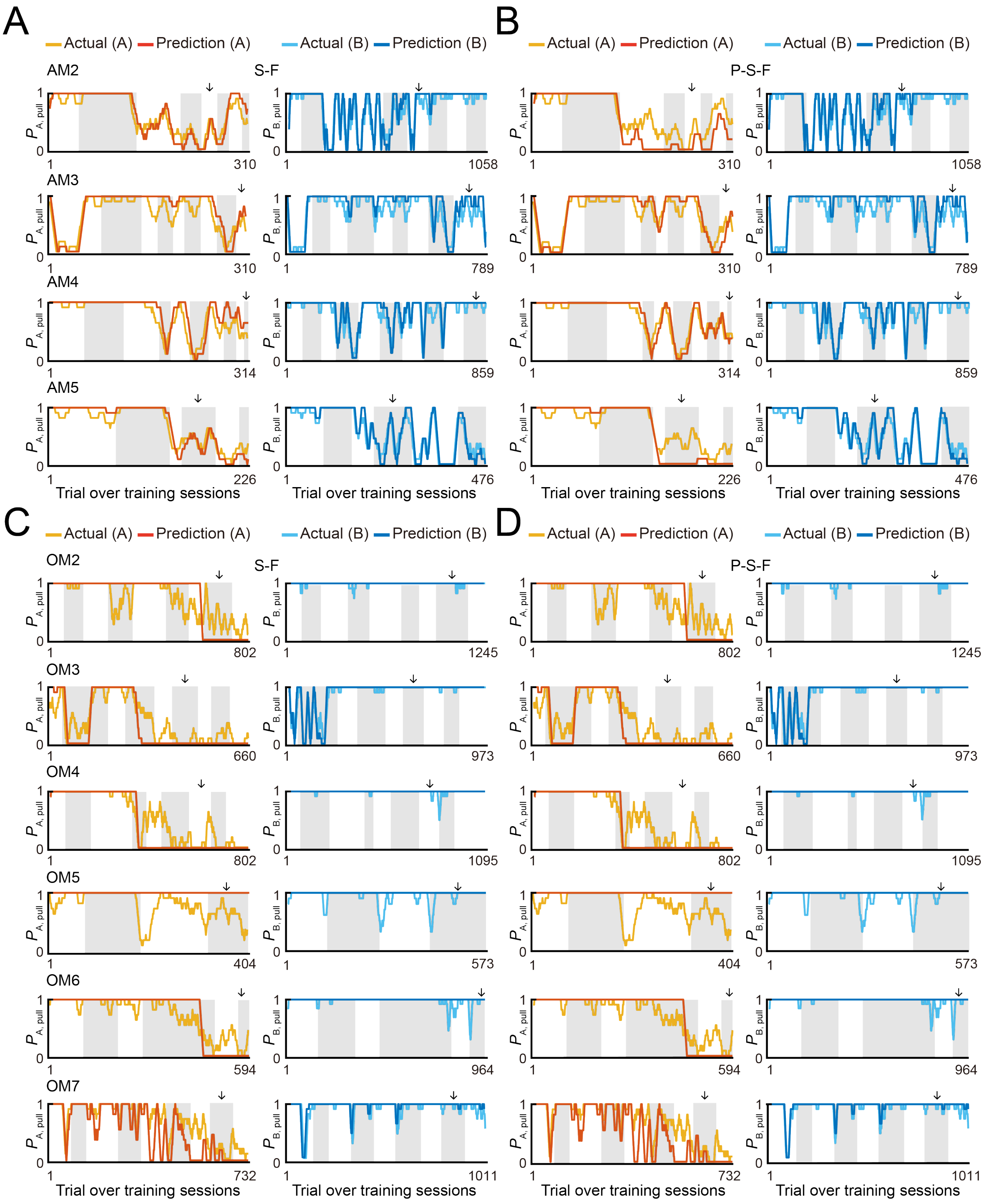


**Figure S1. Predicted pull-choice probability across training sessions.**

**(A, B)** Time course of predicted pull-choice probabilities with the S-F model (A) and P-S-F model (B) in tone A (left, red) and tone B (right, blue) trials for four mice (AM2–AM5) in the air-puff task. The 10-trial moving averages of the actual pull choice in tone A (yellow) and tone B (cyan) trials are overlaid. Even sessions are shaded. Arrows indicate the second threshold sessions.

**(C, D)** Time course of predicted pull-choice probabilities with the S-F model (C) and P-S-F model (D) in tone A (left, red) and tone B (right, blue) trials for six mice (OM2–OM7) in the omission task. The 10-trial moving averages of the actual pull choice in tone A (yellow) and tone B (cyan) trials are overlaid. Even sessions are shaded. Arrows indicate the second threshold sessions.

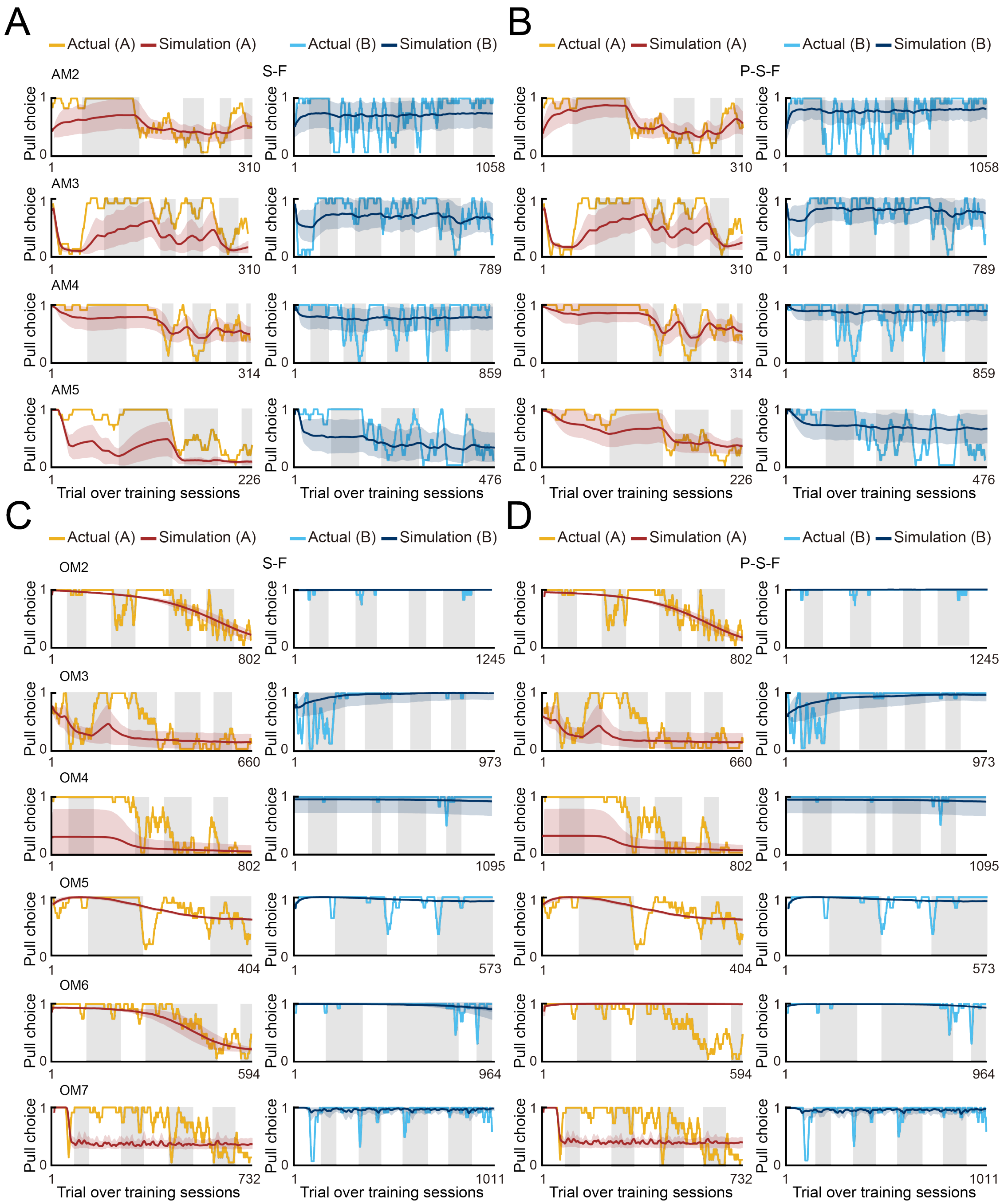


**Figure S2. Simulated pull-choice behavior across training sessions.**
**(A, B)** Simulated pull-choice behaviors with the S-F model (A) and P-S-F model (B) in tone A (left, red) and tone B (right, blue) trials for the AM2–AM5 mice in the air-puff task. For each mouse, simulation was repeated 1000 times and the choice behavior was averaged in each trial (0–1). The 10-trial moving averages of the simulated pull choice in tone A (dark red) and tone B (dark blue) trials are shown. The colored shading indicates ±SD across 1000 simulations. The yellow and cyan traces represent the 10-trial moving averages of the actual pull choices in tone A and tone B trials, respectively. Even sessions are shaded.
**(C, D)** Representative simulated pull-choice behaviors with the S-F model (C) and P-S-F (D) model in tone A (left, red) and tone B (right, blue) trials for the OM2–OM7 mice in the omission task. For each mouse, simulation was repeated 1000 times and the choice behavior was averaged in each trial (0–1). The 10-trial moving averages of the simulated pull choice in tone A (dark red) and tone B (dark blue) trials are shown. The colored shading indicates ±SD across 1000 simulations. The yellow and cyan traces represent the 10-trial moving averages of the actual pull choices in tone A and tone B trials, respectively. Even sessions are shaded.

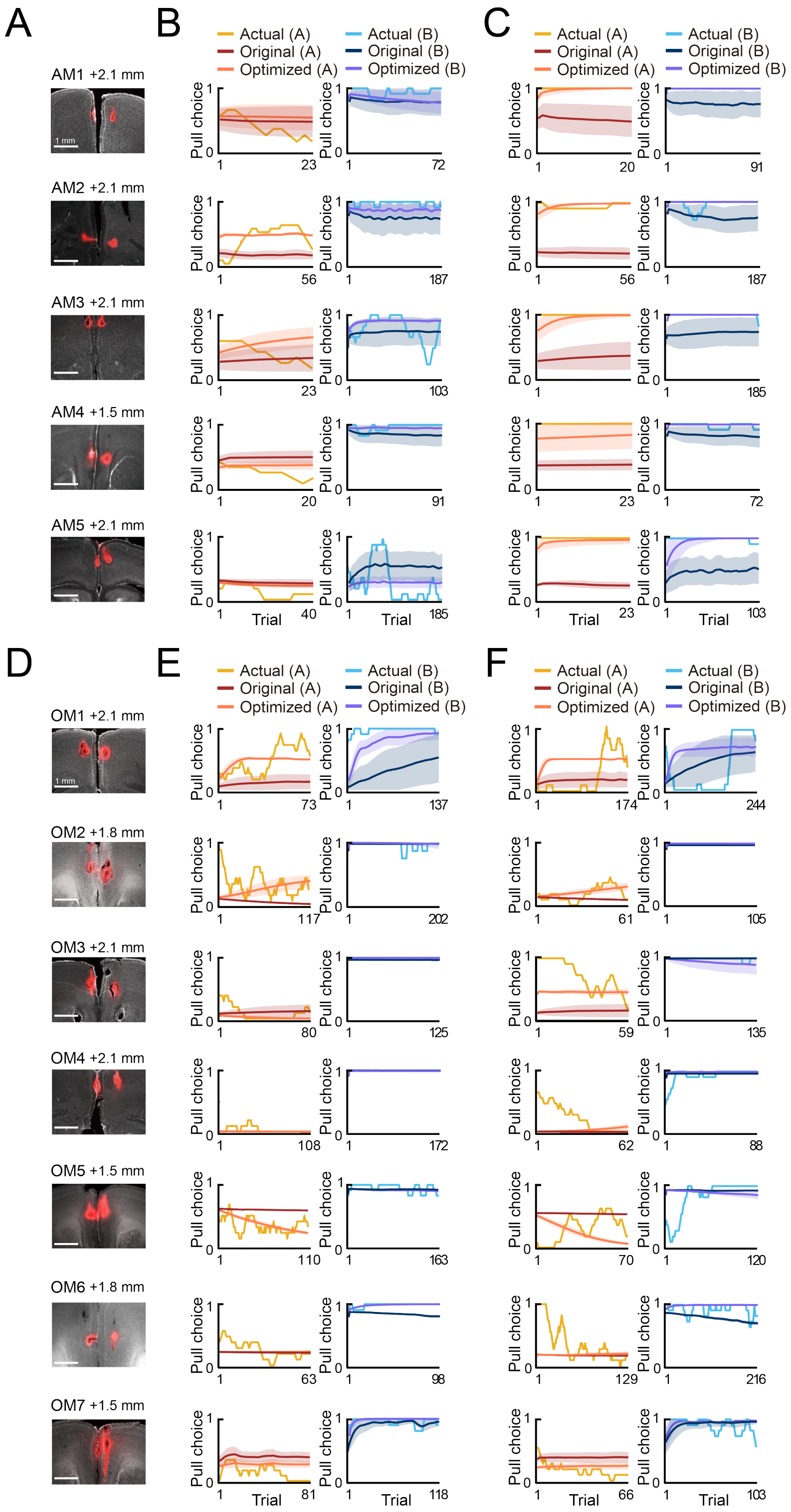


**Figure S3. Injection sites and simulated pull-choice behaviors in the ACSF and muscimol sessions.**
**(A)** Injection sites in the mPFC of each of the mouse AM1–AM5 that performed the air-puff task. NeuroTrace CM-Dil tissue-labeling paste was injected to confirm the injection site after ACSF and muscimol sessions. The number at the upper right of each image indicates the distance from the bregma to the slice along the anterior-posterior axis. **(B, C)** Simulated pull-choice behaviors in the P-S-F model with the best parameter set for the training sessions in the ACSF session (B) and muscimol session (C) in the air-puff task. The yellow and cyan traces represent the actual pull choices. For each mouse, simulation was repeated 1000 times and the choice behavior was averaged in each trial (0–1). The colored shading indicates ±SD across 1000 simulations. The dark red and dark blue traces represent the 10-trial moving averages of pull-choice simulated with the original parameters, whereas the orange and purple traces represent the 10-trial moving average of pull-choice simulated with the optimized parameters. **(D)** Injection sites in the mPFC of each of the OM1–OM7 mice that performed the omission task. NeuroTrace CM-Dil tissue-labeling paste was injected to confirm the injection site after ACSF and muscimol sessions. The number at the upper right of each image indicates the distance from the bregma to the slice along the anterior-posterior axis. **(E, F)** Simulated pull-choice behaviors in the S-F model with the best parameter set for the training sessions in the ACSF session (E) and muscimol session (F) in the omission task. The yellow and cyan traces represent the actual pull choices. For each mouse, simulation was repeated 1000 times and the choice behavior was averaged in each trial (0–1). The colored shading indicates ±SD across 1000 simulations. The dark red and dark blue traces represent the 10-trial moving averages of pull-choice simulated with the original parameters, whereas the orange and purple traces represent the 10-trial moving average of pull-choice simulated with the optimized parameters.

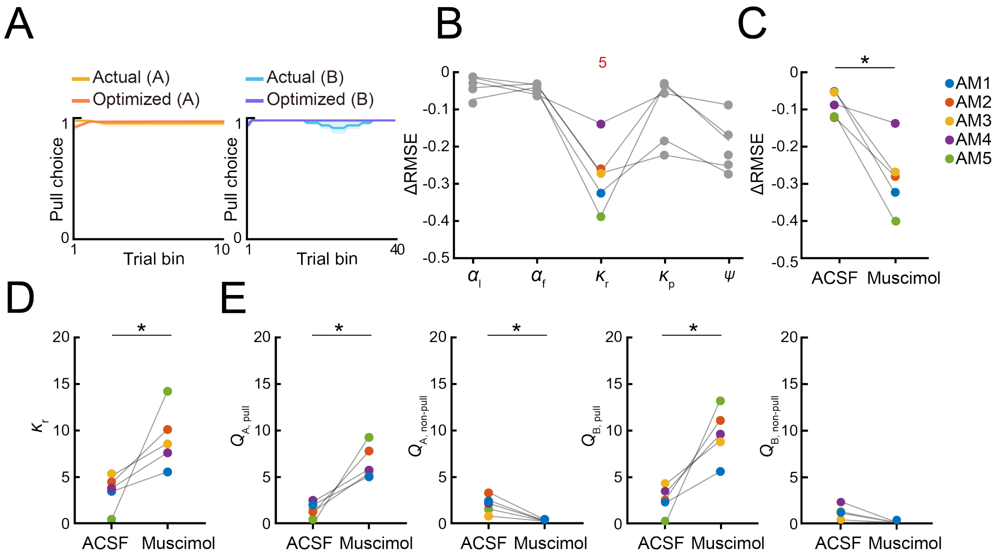


**Figure S4. Simulation of the pull-choice behavior in the muscimol session in the air-puff task after optimization of the parameters including the initial *Q*-values.**

**(A)** Simulated pull-choice behaviors in the P-S-F model with the parameters obtained after the parameter optimization that also included the initial *Q*-values in tone A (left) and tone B (right) trials in the muscimol session in the air-puff task. The yellow and cyan traces represent the actual pull choices, whereas the orange and purple traces represent the simulated pull-choice. For each mouse, the simulation was repeated 1000 times and the choice behavior was averaged in each trial (0–1). To average the choice behavior across the mice with different trial numbers, the trials in the session were divided into 10 bins in tone A trials and 40 bins in tone B trials for each mouse, and the choice behavior was averaged within each bin. Then, for each bin, the choice behavior was averaged over the mice. The shading indicates ±SEM across the animals (n = 5). **(B)** Improvement of the simulation of the choice behavior with optimization of each parameter that also changed the initial *Q*-values in the muscimol sessions in the air-puff task. Each line indicates a different animal (AM1–AM5). Each colored dot indicates the parameter that minimized the error between the actual and simulated choice behaviors (ΔRMSE) for each mouse. The red number indicates the number of colored dots for the corresponding parameter. **(C)** The change in the RMSE (ΔRMSE) resulting from optimization of parameters that also included the initial *Q*-values when simulating the behaviors in the ACSF and muscimol sessions. Each color indicates a different mouse. *p = 0.0195, paired *t* test. **(D)** κr for optimization in the training sessions and for optimization that also included the initial *Q*-values in the muscimol session. Each color indicates a different animal (AM1–AM5). *p = 0.0215, paired *t* test (n = 5). **(E)** $Q_{A,\ pull}$, $Q_{A,\ non\text{–}pull}$, $Q_{B,\ pull}$, and $Q_{B,\ non\text{–}pull}$ estimated with the optimized parameters that also included the initial *Q*-values in the ACSF and muscimol sessions. Each colored line represents an individual mouse (AM1–AM5). Each dot indicates the average within each session. $Q_{A,\ pull}$, *p = 0.0254; $Q_{A,\ non\text{–}pull}$, *p = 0.0102; $Q_{B,\ pull}$, *p = 0.0128; $Q_{B,\ non\text{–}pull}$, p = 0.1885, paired *t* test (n = 5).

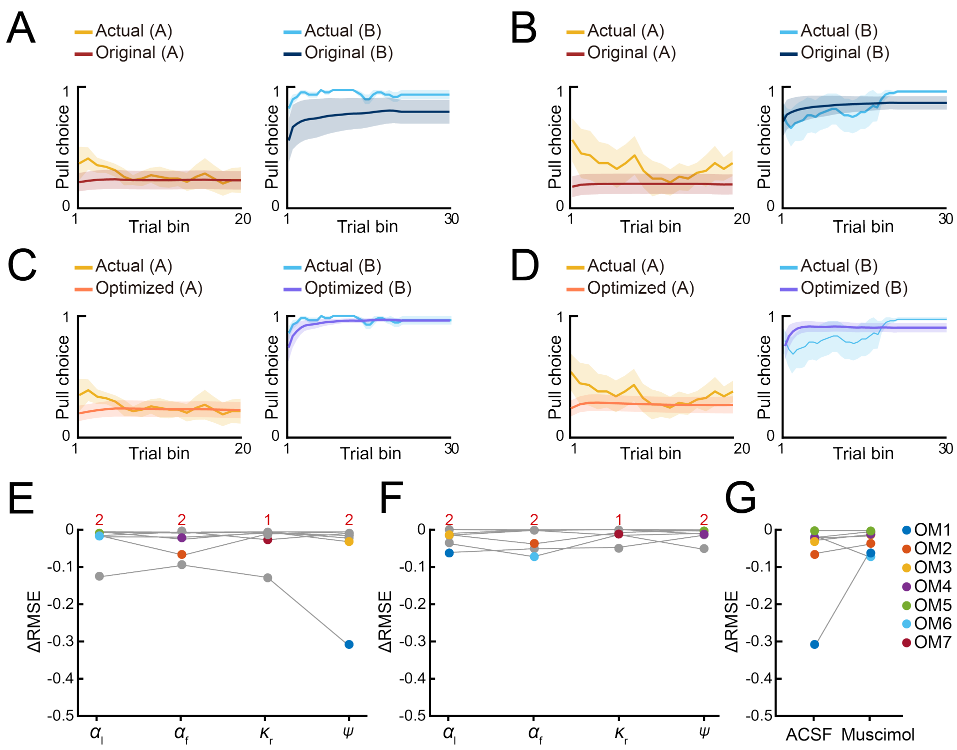

**Figure S5. Parameter optimization to explain the pull-choice probability under inhibition of mPFC in the omission task.**
**(A, B)** Simulated pull-choice behaviors in the S-F model with the best parameter set for the training sessions in the ACSF session (A) and muscimol session (B) in the omission task. For each mouse, the simulation was repeated 1000 times and the choice behavior was averaged in each trial (0–1). To average the choice behavior across the mice with different trial numbers, the trials in the session were divided into 20 bins in tone A trials and 30 bins in tone B trials for each mouse, and the choice behavior was averaged within each bin. Then, for each bin, the choice behavior was averaged over the mice. The yellow and cyan traces represent the actual pull choices, whereas the dark red and dark blue traces represent the pull-choice simulated with the original parameters. The shading indicates ±SEM across the animals (n = 7). **(C, D)** Simulated pull-choice behaviors in the S-F model with the parameters after the parameter optimization in the ACSF session (C) and muscimol session (D) in the omission task. The yellow and cyan traces represent the actual pull choices, whereas the orange and purple traces represent the pull-choice simulated with the parameters after parameter optimization. The shading indicates ±SEM across the animals (n = 7). **(E, F)** Improvement of the simulation of choice behavior with optimization of each parameter in the ACSF session (E) and muscimol session (F) in the omission task. Each line indicates a different animal (OM1–OM7). Each colored dot indicates the parameter that minimized the error between the actual and simulated choice behaviors (ΔRMSE) for each mouse. Red numbers indicate the number of colored dots for the corresponding parameter. **(G)** The change in RMSE (ΔRMSE) resulting from parameter optimization in the simulation of the behaviors in the ACSF and muscimol sessions. Each colored line indicates an individual mouse (OM1–OM7). p = 0.3557, paired *t* test.
